# Optimization of liver graft function using poly-pharmacological drug cocktail CEPT in a simulated transplant model

**DOI:** 10.1101/2024.02.02.578568

**Authors:** Anil Kharga, Mohammadreza Mojoudi, Huyun Chen, McLean S. Taggart, Antonia T. Dinicu, Ozge S. Ozgur, Basak Uygun, Mehmet Toner, Shannon N. Tessier, Heidi Yeh, James F. Markmann, Alban Longchamp, Korkut Uygun

**Author notes:** **Corresponding author:** Korkut Uygun, PhD.

## Abstract

**Background:** The number of patients in need of a liver transplant far exceeds the supply of available organs. This imbalance could be dramatically reduced should the donor organ pool be expanded by rendering marginal cases transplantable rather than discarded. The poly-pharmacological drug cocktail CEPT (Chroman-1, Emricasan, Polyamine, and Trans-ISRIB (integrated stress inhibitor)) has been found to improve the in-vitro viability of human pluripotent stem cells (hPSCs) following cryopreservation. It is worth exploring CEPT’s ability to inhibit various apoptotic pathways and preserve cellular function for potentially mitigating warm ischemic stress of the anhepatic phase of graft implantation and promoting more rapid graft recovery following reperfusion with continuous treatment.

**Methods:** Rat livers without warm ischemia and CEPT supplementation are the healthy control: fresh (n=3) group. Room-temperature warm ischemia was used to replicate the anhepatic phase of graft implantation in the control (n=6) group and the experimental CEPT group (n=6) without and with CEPT supplementation, respectively. Transplantation was modeled by ex-vivo reperfusion at 37°C for six hours with William’s E-based hepatocyte culture media and with CEPT supplementation in the CEPT group.

**Results:** Livers treated with CEPT during warm ischemia and subsequent reperfusion have improved hepatocellular function as indicated by increased oxygen O_2_ utilization, stable pH, and improved cholangiocyte function indicated by the increased hourly rate of bile production. Furthermore, resistance, an endothelial injury marker, and caspase 3/7, an apoptotic marker, were lower.

**Conclusion:** To improve the utilization of available donor livers, different stages of the organ transplantation process can be optimized. The anhepatic phase, which includes the period from the removal of the native liver from the recipient to the reperfusion of the donor’s graft liver through the portal vein during graft implantation, can be targeted using CEPT for mitigating warm ischemia-induced injury that occurs during vascular anastomosis. **(S1 Fig: Graphical abstract)**

## Introduction

Liver transplantation is a standard procedure with high survival rates. However, 15% of patients die while awaiting a viable liver [1]. Typically, grafts are derived from donors who have suffered cardiac death or are brain dead. Each source of graft poses different challenges for transplantation. For example, grafts procured from controlled donation after cardiac death (DCD) suffer warm ischemic damage. Damage to grafts from a brain-dead donor (DBD) is primarily because of sterile inflammation from the cytokine storm, imbalance of the autonomic nervous system, and loss of hypothalamic-pituitary-adrenal axis, which causes electrolyte and hemodynamic imbalance [2]. Although DBD grafts do not generally suffer severe warm ischemic damage like that which occurs in a DCD, both types of grafts suffer from cold ischemic damage during transportation and warm ischemic damage during the anhepatic phase of graft implantation in the recipient. Thus, to improve utilization of liver grafts, it is crucial to explore ways to minimize ischemic damage at various stages of transplantation. In the current work, we focused on utilizing drug intervention during the anhepatic phase of graft implantation.

The anhepatic phase of graft implantation is the duration from the removal of the native liver of the recipient to reperfusion of the donor’s graft liver through the portal vein in the recipient [3]. Liver core temperature rises rapidly towards room temperature during the performance of vascular anastomosis [4]. Increased temperature increases the graft’s metabolic demand [5]; as a result, the graft suffers from warm ischemic damage after the liver is removed from storage on ice at the end of back-benching during vascular anastomosis. Furthermore, anhepatic phases over 100 minutes have a higher incidence of graft dysfunction and worse 1-year patient survival [3], and increased implantation time is associated with poor transplant outcomes due to a rise in the graft temperature [6]. Preclinical animal studies to randomized clinical trials (RCT) have used drugs to precondition the graft before implantation and have treated the recipient to help the graft recover faster upon reperfusion [7–10]. Using a drug to target the anhepatic phase of graft implantation and treating the recipient with a drug for faster recovery could be a more straightforward and practical approach.

A new poly pharmacological drug cocktail-**CEPT**: **C**hroman 1, **E**mricasan, **P**olyamines, and **T**rans-ISRIB (integrated stress inhibitor) has been shown to enhance cell survival of genetically stable human pluripotent stem cells (hPSCs) and their differentiated progeny by simultaneously blocking several stress and apoptotic pathways that otherwise would have compromised cell structure and function [11]. **Chroman 1** is a highly potent ROCK (Rho-associated coiled-coil forming protein serine/ threonine kinase) pathway inhibitor [12], which improves cell survival by blocking cell contraction through decreased phosphorylation of the myosin light chain, resulting in inhibition of action-myosin contraction, ultimately preventing apoptosis [13]. **Emricasan**, a pan-caspase inhibitor, efficiently blocks caspase-3 activation alone and, when combined with Chroman 1, has proved superior to single compound use [11]. **Trans-ISRIB**, a selective inhibitor of the integrated stress response (ISR), makes cells resistant by preventing stress granule formation and restoring the messenger ribonucleic acid (mRNA) translation [14]. **Polyamines** are positively charged metabolites involved in cellular functions such as transcription, translation, cell cycle, and stress protection [15]. We hypothesize that by targeting various apoptotic pathways, CEPT may minimize Ischemic Reperfusion Injury (IRI) and improve graft recovery post-reperfusion in the recipient.

Recent clinical trials have demonstrated ex-vivo machine perfusion’s efficacy in reconditioning ischemic damaged grafts and assessing the viability of DCD, elderly, and steatotic grafts during the preservation period [16]. Machine perfusion can potentially recondition grafts by mitigating IRI [17]. Normothermic Machine Perfusion (NMP) simulates near-physiological conditions to maintain normal metabolic functions, allowing graft assessment using the liver’s hemodynamic, biochemical, and synthetic functions and bile analysis [18]. In addition, machine perfusion can be applied to study drug pharmacokinetics to predict newly developed drugs’ absorption, distribution, metabolism, excretion profile, and toxicological behavior [19]. In the current work, we use NMP to simulate the transplantation process on the pump for recovery of the rat liver graft using the drug cocktail-CEPT and simultaneous assessment of graft function.

## Methods

### Ethics statement

This study complies with ethical regulations, and the experimental protocol no: 2011N000111 was approved by the Institutional Animal Care and Use Committee (IACUC) of Massachusetts General Hospital, Boston, MA, USA.

### Rationale for the study design

In a 45 min Warm Ischemia Time (WIT) donation after brain death (DBD) porcine model of liver transplantation, a combination of drugs was used for flushing and another combination of drugs was used in the recipient, which decreased the primary non-function of the graft, improved liver function and increased survival [9]. In a recent randomized control trial, following static cold storage, donor’s livers were infused ex-situ with epoprostenol via portal vein; recipients received oral α-tocopherol and melatonin before anesthesia and during the anhepatic and reperfusion phase received intravenous drugs-antithrombin III, infliximab, apotransferrin, recombinant erythropoietin-β, C1-inhibitor, and glutathione [7]. In adult living donor liver transplantation clinical trials, intraoperative dexmedetomidine infusion decreased IRI by suppressing intercellular adhesion molecule-1 (ICAM-1) [8]. In a rat orthotropic liver transplant model, Treprostinil, a prostacyclin (PGI 2) analog, was given to donor animals 24 hours before hepatectomy. The recipient animal received a similar treatment until the time of sacrifice, and this drug intervention ameliorated IRI [10].

Based on these findings, we used NMP technology to simulate the transplantation of rat liver on a pump to explore a new drug cocktail-CEPT efficacy in helping the liver graft, which always suffers ischemic damage due to a gradual rewarming towards room temperature during the anhepatic phase of graft implantation in the recipient, by flushing the graft with the drug before the simulated graft implantation phase and simulating treatment of recipient post-reperfusion using NMP pump.

### Study design

The study design is illustrated in **Fig 1. A**. and it is described below:

**1) CEPT** group - experimental group: (n=6): The rat liver was procured by flushing with room temperature Lactated Ringer’s (LR) for 15 minutes with CEPT. Procurement was followed by 60 minutes of warm ischemia at room temperature (21°C). During WIT, the liver was placed in 50 ml of LR, supplemented with CEPT, mimicking the recipient’s anhepatic phase of graft implantation. During this phase, the drug is present in the graft vasculature. It is aimed to decrease apoptosis during warm ischemia, then 6 hours of NMP with CEPT-supplemented perfusate (500ml), mimicking post-reperfusion phase in the recipient with the drug treatment to improve graft recovery from the ischemic damage suffered during procurement and anhepatic phase of graft implantation.
**2) Control** group (n=6): The rat liver was procured by flushing with room temperature LR for 15 minutes without CEPT. The liver was then stored in 50 mL of LR without CEPT for 60 minutes at room temperature (21°C) to mimic WIT. Following 60 minutes, the liver underwent 6 hours of NMP with 500 mL of perfusate without CEPT.
**3) Fresh** group-healthy control: (n=3) The rat liver was procured by flushing with room temperature LR without CEPT for 15 minutes without the drug (CEPT), then it was immediately transferred to the NMP pump with perfusate (500ml) not supplemented with CEPT. The fresh liver did not experience 60 minutes of WIT at room temperature (21°C), unlike the CEPT and control groups.

**Fig 1:**
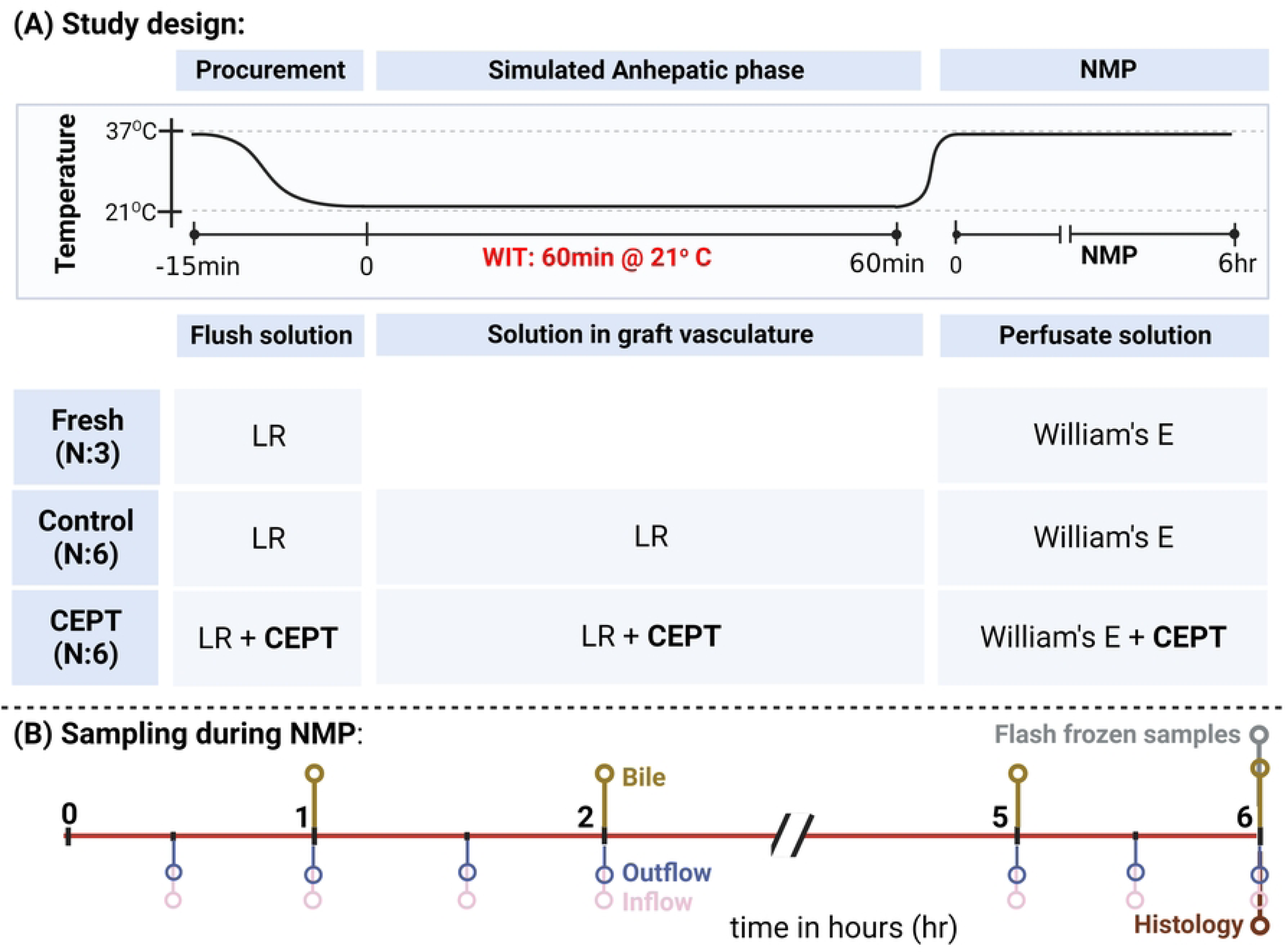
Study design and sampling timeline during NMP: (A) Study design: Rat liver was flushed with LR with/ without drug cocktail (CEPT) during procurement. Fresh group livers were flushed without the drug cocktail and were transferred to the NMP pump immediately; CEPT and control group livers were kept in LR solution with/without the drug cocktail, respectively, for 60 minutes at room temperature (21oC) to induce a low level of warm ischemia of anhepatic phase of graft implantation before transferring to the NMP pump to simulate transplantation with/ without drug cocktail in perfusate respectively. (B) Sampling during NMP was conducted half hourly for perfusate from inflow and outflow ports, bile was collected hourly, and tissue biopsy for flash frozen samples and histology was taken at the end of NMP. CEPT: Chroman-1, Emricasan, Polyamine, and Trans-ISRIB (integrated stress inhibitor); WIT: Warm Ischemia Time; LR: Lactated Ringer’s, NMP: Normothermic Machine Perfusion.

### Rat hepatectomy

Adult male Lewis rats (weighing 300-350 g) were obtained from Charles River Laboratories, Wilmington, MA, USA. The animal study was performed according to the National Research Council guidelines and the Institutional Animal Care and Use Committee (IACUC) protocols at Massachusetts General Hospital (protocol number: 2011N000111). Male Lewis rats were housed socially in temperature (70 °F ± 2 °F) and humidity (30–70%) controlled environments within pathogen-free high-efficiency particulate air (HEPA)-filtered ventilated cages, alternating 12-hour light/dark cycles. Animals were provided sterilized standard rat chow and water ad libitum. Fifteen rats underwent total hepatectomy. Male Lewis rats were first anesthetized by inhalation of 5% isoflurane in an induction chamber. During the surgical procedure, anesthesia was administrated using a tabletop anesthesia apparatus connected to a standard rodent system. Rats were considered adequately anesthetized when a muscular contraction was absent following a toe pinch. After anesthesia induction, the rat was placed on a surgical table in a supine position and covered with sterile surgical drapes. The abdomen was opened with a transverse abdominal incision 1 cm below the lower end of the sternum. Ligaments and adhesions at the superior and posterior aspects of the liver were divided to free the liver from its surroundings. The rats’ bile duct was cannulated with 27 G tubing and secured in place with 6-0 silk thread.

After which, portal vein tributaries-splenic and gastric veins and hepatic artery were ligated with 6-0 silk. Heparin (1 unit/g of rat weight) was injected into the infra-hepatic inferior vena cava (IVC). The portal vein (PV) was cannulated with an 18-G Angio catheter (green colored), and the infra-hepatic IVC was transected immediately to flush the liver and minimize backward building pressure from the flushing fluid. The animal was euthanized by exsanguination of blood from transected IVC. The liver was then flushed with 50 mL room temperature (21°C) Lactated Ringer’s (LR) with supplementation of 1 mL heparin in the 50 mL LR through the PV. The PV cannula was secured with 6-0 silk thread, and the liver was freed from its remaining attachments to the diaphragm, retroperitoneum, and intestine. The supra-hepatic IVC was transected, and the liver was removed from the body and weighed immediately to get the initial weight used for calculating edema, oxygen consumption, portal vascular resistance, and hourly rate of bile production.

### Perfusate composition

Ex-vivo NMP was performed with a sterilized and oxygen carrier free perfusate. 1L of perfusate was composed of 950 mL of William’s E hepatocyte culture media (Sigma Aldrich, Burlington, MA, USA) to supply the graft with amino acids and nutrition during perfusion, 20 g of bovine serum albumin (BSA) (Sigma Aldrich, Burlington, MA, USA) for oncotic pressure required for balancing portal hydrostatic pressure to prevent tissue edema, 20 g of 35,000 kilo-Dalton polyethylene glycol (PEG) (Sigma Aldrich, Burlington, MA, USA), 20 mg of dexamethasone (Sigma Aldrich, Burlington, MA, USA), 2 mL (1000 unit/ml) of heparin (MGH Pharmacy, Boston, MA, USA), 1 mL (100 units/ml) of regular insulin (MGH Pharmacy, Boston, MA, USA), 10 mL (5000 units/ml of penicillin and 5 mg/ml of streptomycin) penicillin-streptomycin (Thermo Fisher Scientific, Point Street, CA, USA), and 2.2 g of sodium bicarbonate to obtain a pH of 7.4. On the day of the experiment, 5 mL of L-glutamine (Sigma Aldrich, Burlington, MA, USA) was added to 500 mL perfusate prior to perfusion since L-glutamine degrades after 24 hours of perfusate preparation. The CEPT group perfusate was supplemented with the drug cocktail-CEPT: 50 nM Chroman 1 (MedChemExpress, Monmouth Junction, NJ, USA), 5 uM Emricasan (MedChemExpress, Monmouth Junction NJ, USA), 0.7 uM trans-ISRIB (Tocris Bioscience, Emeryville, CA, USA), 40 ng/mL putrescine (Sigma Aldrich, Burlington, MA, USA), 4.5 ng/mL spermidine (Sigma Aldrich, Burlington, MA, USA), and 8 ng/mL spermine (Sigma Aldrich, Burlington, MA, USA), all suspended in dimethyl sulfoxide (DMSO) (Sigma Aldrich, Burlington, MA, USA). Putrescine, spermidine, and spermine form the polyamine portion of the CEPT.

### Normothermic machine perfusion (NMP)

**Fig 2** illustrates the NMP setup. 500 mL of perfusate was circulated using a peristaltic roller pump system for a 6-hour duration at 37°C at a constant flow rate of 30 mL/min through a membranous oxygenator connected to a 95% O_2_ and 5% CO_2_ gas cylinder (Airgas, Radnor, PA, USA). The PV inflow pressure was kept below 12 mmHg. Perfusate entered the PV and exited freely from the supra-hepatic inferior venacava (IVC) and infra-hepatic IVC. The perfusion circuit consisted of a perfusion chamber: tissue bath (Radnoti, Covina, CA, USA), Masterflex peristaltic pumps (Cole Parmer, Vernon Hill, IL, USA), a membrane oxygenator (Radnoti, Covina, CA, USA), a heat exchanger (Radnoti, Covina, CA, USA), a bubble trap (Radnoti, Covina, CA, USA). The liver temperature was regulated by a water bath (Lauda, Brinkmann, Westbury, NY, USA) and constantly monitored. During perfusion, the pH was monitored without the supplementation of bicarbonate to assess the ability of the graft to maintain near physiological pH on its own.

**Fig 2:**
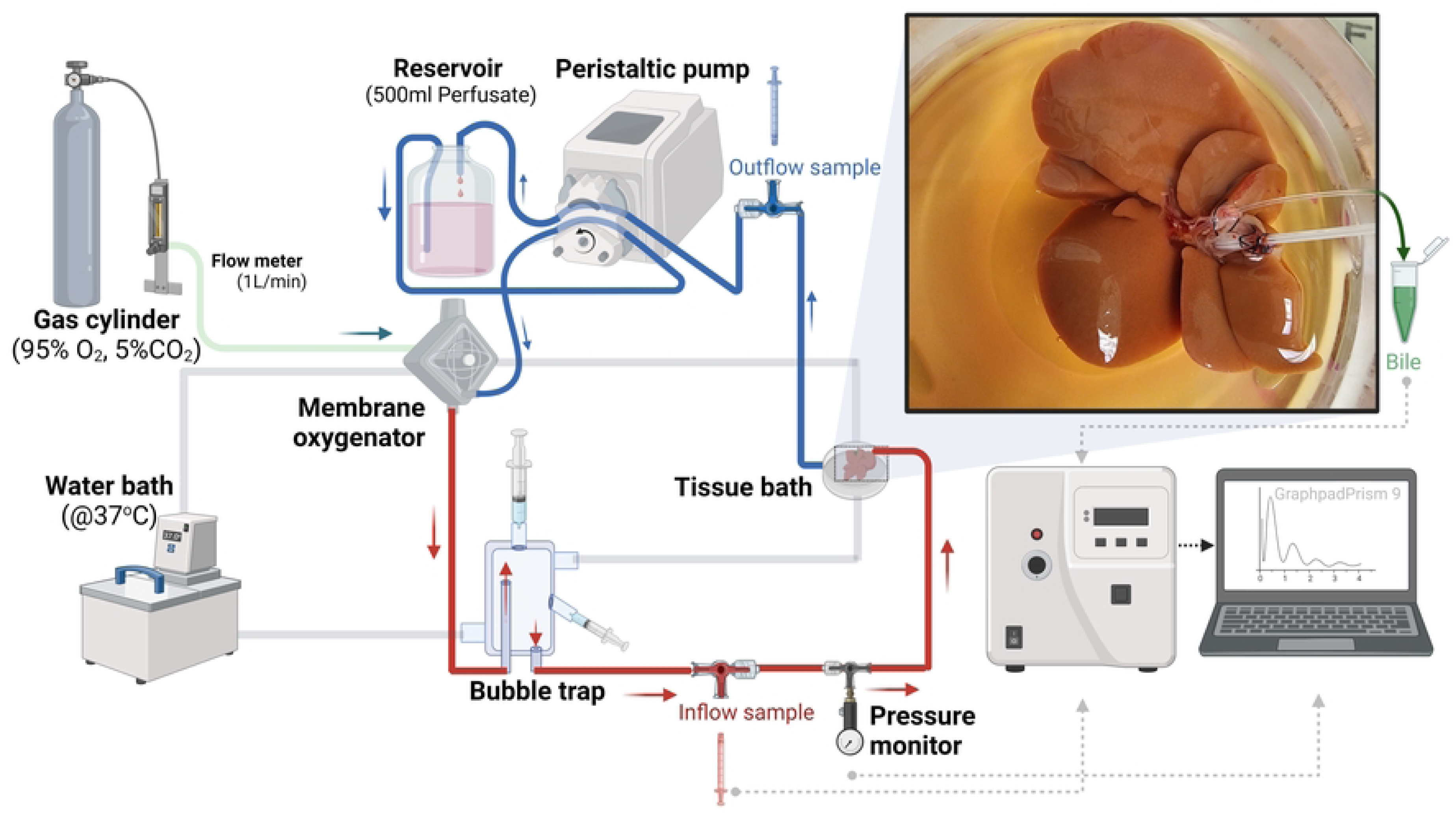
Normothermic Machine Perfusion (NMP) of rat liver. The peristaltic pump pushes perfusate from the reservoir through the membrane oxygenator to raise partial pressure (pO_2_) above 500 mmHg and removes CO_2_ from perfusate. The bubble trap prevents air emboli and acts as a compliance chamber, along with 50ml Syringes to dampen pulsation from the pump. The water bath heats water to 37oC and pumps warm water through the jacketed structure of the oxygenator, tissue bath, and bubble trap to keep perfusate at body temperature (37oC). Samples from inflow and outflow ports were collected every half an hour for analysis, and pressure values were recorded using a pressure transducer attached to the inflow line. Bile was collected every hour for analysis, and the volume produced was recorded.

### Sample collection

The sample collection timeline is illustrated in **Fig 1. B**. A proximal port was used for the inflow sample collection, and the outflow sample was collected directly from infra-hepatic IVC. The baseline sample was collected at 15 minutes when the liver had reached the maximum flow rate set (30 ml/min), and bile was collected hourly under the mineral oil to prevent pCO_2_ from equilibrating with the atmosphere and have accurate readings for bile pH and bile HCO_3_. After the baseline sampling, inflow and outflow perfusates were analyzed using the Siemens RapidPoint 405 Blood Gas Analyzer (Siemens Healthineers, Malvern, PA, USA). Additional perfusion samples were collected and stored every half an hour for subsequent analysis. Liver biopsies and flash-frozen samples were collected at the end of NMP. The rat liver tissue samples collected were fixed in 10% formalin, embedded in paraffin, and sectioned.

### Assessment of graft function

Liver hepatocellular functions, injury markers, and cholangiocyte functions were assessed on the pump using NMP after inducing 60 minutes of warm ischemic damage at room temperature to mimic the anhepatic phase of graft implantation.

#### 1) Metabolic function

The metabolic function was assessed with oxygen (O_2_) uptake rate and the ability to maintain near physiological pH without the need for bicarbonate (HCO_3_) supplementation. Oxygen uptake or consumption rate was defined as inflow partial pressure (pO_2_ in) minus outflow pO_2_ (pO_2_ out) times flow rate (30ml/min) per gram (g) of liver weight. Oxygen uptake in µL/min/g = (pO_2_ in – pO_2_ out) * Flow rate * 0.314/ weight.

#### 2) Injury markers

Liver graft injury was accessed using portal vascular resistance, ineffective glucose metabolism from glucose level, potassium (K), lactate, and liver transaminase: alanine transaminase (ALT) and aspartate transaminase (AST) from the portal vein’s inflow line and caspase 3/7 activity at the end of perfusion from the flash-frozen samples as a marker of apoptosis. In addition, edema compared to initial weight at procurement due to WIT for 60 minutes at room temperature before the start of NMP for simulating anhepatic phase of graft implantation and weight changes at the end of NMP compared to initial weight at time of procurement were accessed for cellular and interstitial edema and its resolution at the end of NMP, respectively. Portal vascular resistance was defined as pressure divided by flow rate per gram (g) of liver weight as an indirect marker of liver sinusoidal endothelial cell edema.

#### 3) Cholangiocellular function

The volume of bile produced every hour was recorded as hourly the rate of bile production, and it was analyzed for biochemistry to calculate bile parameters delta values. Bile pH delta, bicarbonate (HCO_3_) delta, and K delta were calculated by subtracting the perfusate inflow values from the bile values at the hourly time point.

#### 4) Histological analysis

Hematoxylin-eosin (H&E) and Terminal deoxynucleotide transferase dUTP nick end labeling (TUNEL) stains were utilized to segment and stain fixed liver tissue samples collected at the end of 6 hours of perfusion (Specialized Histopathology Services Core, MGH, Charlestown, MA, USA).

#### 5) Storage solution and effluent drainage analysis

CEPT and Control groups’ livers were kept in 50 mL of LR solution at room temperature for 60 minutes to simulate the gradual rewarming phase of the anhepatic phase of graft implantation in the recipient. This storage solution (50 mL LR) was analyzed for biochemistry and transaminase level. Effluent drainage collected from IVC after flushing with 3 mL of room temperature LR over 1 minute just before transferring the liver to the NMP pump was analyzed for biochemistry panels: pH, Lactate, Glucose, K, and transaminase: ALT & AST. Point of Care (POC) machine could not detect pH lower than 6.5 and glucose lower than 20 mg/dL. These missing data points were replaced with the lowest measured value of 6.5 for pH and 20 mg/dL for glucose in the graph and for data analysis.

### Caspase 3/7 Activity Assay

For assessments of caspase 3/7 activity, flash-frozen biopsy samples were collected at the end of each perfusion. For this, 300-500 mg of livers were flash-frozen in liquid nitrogen for preservation at –80°C. On the day of measurement, frozen liver samples were added to ice-cold PBS at 50 mg/mL and processed for tissue homogenization using the gentleMACS Dissociator (n = 3 per group) (Miltenyi Biotec, Bergisch Gladbach, Germany). Then, an equal volume of room-temperature PBS was added to the lysate to dilute it to 25 mg/mL. In a white flat bottom 96-well assay plate, 100 uL of the diluted lysate was added to each well along with an equal volume of Caspase Glo 3/7 assay reagent (Promega, Madison, Wisconsin, USA), according to the manufacturer’s directions. The plate was then placed on a plate shaker set at 250 rpm for 30 minutes at room temperature while protected from light. After incubation, luminescence was read from the top of the well using the Synergy 2 microplate reader (Biotek, Winooski, VT, USA).

### Statistical Analysis

Data were managed using Microsoft Excel 365. Statistical analyses and graphs were generated with Prism 9.5.1 (GraphPad Software Inc., La Jolla, CA, USA). Ordinary one-way ANOVA was used for one-factor comparison with Tukey’s post-hoc test for multiple comparisons. The significance level was set for p-value < 0.05. Difference among groups at each time point of sample collection was determined through repeated measures, two-way ANOVA followed by Tukey’s post-hoc test for multiple comparisons. Simple linear regression was used to show linear trends over time, such as for liver transaminase level (AST/ALT), and their slopes were compared using ANOVA. For edema developed due to induced warm ischemia (for two groups: CEPT and control only), two-tailed two-sample student’s t-test with Welch’s correction was used. For pH, K, and glucose, pre-perfusion (time: 0) inflow values were recorded when the liver was not placed on the system to show the normal perfusate concentration and pH in the graph, but it was excluded for analysis as the liver was not present for assessment at time: 0.

## Results

### 1) Metabolic function

The CEPT group had a significantly higher oxygen uptake rate (**Fig 3. A**) than the control group (40.37 ± 0.44 vs. 38.00 ± 0.35 μL of O_2_ /min/g respectively, p = 0.0003), and slightly higher than the fresh group (38.81±0.35, p = 0.017). However, control and fresh groups were not different from each other. CEPT group inflow pH (**Fig 3. B**) was significantly higher than the control group (7.326±0.004 vs. 7.283±0.004 respectively, p<0.0001), and was similar to the fresh group (7.318 ± 0.006).

**Fig 3:**
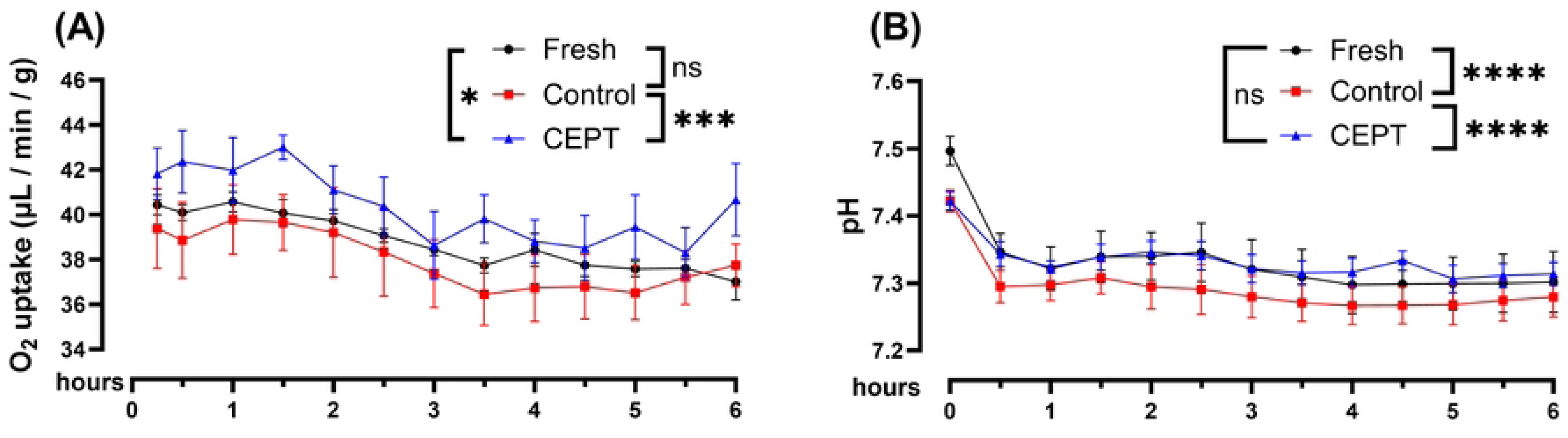
Metabolic function during Normothermic Machine Perfusion (NMP): (A) Oxygen (O_2_) uptake, (B) pH (inflow). O_2_ uptake rate and pH were significantly better in CEPT treated group compared to the control group. Fresh (n=3): healthy control without 60 min warm ischemia time (WIT) at 21oC, control (n=6): with 60 min WIT @21oC but without CEPT, CEPT (n=6): experimental group with 60 min WIT @ 21oC and with CEPT; ANOVA with Tukey’s post-hoc test for multiple comparisons; results are expressed as mean with SEM; pre-perfusion (time: 0) inflow pH values before livers were placed on the pump were excluded for analysis; ns: p > 0.05, * : p ≤ 0.05, ** : p ≤ 0.01, *** : p ≤ 0.001, **** : p ≤ 0.0001.

### 2) Injury markers

Overall, portal vascular resistance (**Fig 4. A**) of CEPT-treated liver was significantly lower than the control (0.01381± 0.0005 vs. 0.01788± 0.0005 mmHg/(ml/min)/g of liver respectively, p <0.0001). However, both CEPT and control groups had portal vascular resistances significantly higher than the fresh group (0.01±0.0005 mmHg/(ml/min)/g of the liver; CEPT vs. fresh and control vs. fresh, both p <0.0001) based on one-way ANOVA. Based on two-way ANOVA, the control group resistance was significantly higher than the fresh liver at 1 hr (0.018±0.003 vs. 0.008±0.001 mmHg/(ml/min)/g, p = 0.04) and 5 hr (0.015±0.002 vs. 0.010±0.001 mmHg/(ml/min)/g, p = 0.03). Glucose levels (**Fig 4. B**) of both CEPT and control groups were similar (227.6± 3.213 vs. 223.1±4.435 mg/dL respectively, p=0.75) and both were significantly higher than the fresh group (198.4±5.378 mg/dL; fresh vs. CEPT p=0.0001 and fresh vs. control p=0.0011). Although the Potassium (K) (**Fig 4. C**) of the CEPT and control groups were not different overall for the entire 6 hours (5.118±0.0099 vs. 5.166±0.0153 mMole/L respectively, p= 0.1150), it is lower in the first half, then reaches a similar level to that of control in the second half for the CEPT group. However, in both groups, K levels are higher than the Fresh group (5.015±0.022 mMole/L; fresh vs. control p<0.0001 and fresh vs. CEPT p= 0.0002). Overall CEPT group lactate (**Fig 4. D**) level was not different from the control group (1.119±0.0354 vs. 1.239±0.065 mMole/L, respectively), but the fresh group’s lactate (1.040±0.062 mMole/L) was significantly lower than the control (p=0.04) based on one-way ANOVA. However, from two-way ANOVA, the CEPT group lactate was lower than the control at 0.5 hr of perfusion (1.310±0.145 vs. 1.597±0.156 mMole/L respectively, p=0.027). Liver transaminase (ALT/ AST) (**Fig 4. E** and **Fig 4. F**) levels raised over time, showing a linear trend with time on running simple linear regression (ALT: R^2^ values for fresh, control, and CEPT: 0.55, 0.31, 0.35, respectively; AST: R2 values for fresh, control, and CEPT: 0.57, 0.32, 0.39, respectively); however, their slopes were not significantly different (ALT slopes for fresh, control, and CEPT: 15.10±3.43, 19.41±5.17, 25.66±6.16, respectively, p=0.45 and AST slopes for fresh, control, and CEPT: 17.41±3.77, 20.79±5.44, 25.89±5.77, respectively, p=0.53). Caspase 3/7 activity (**Fig 4. G**), a marker for apoptosis, measured from flash-frozen samples collected at the end of NMP, was significantly lower in the CEPT group compared to both the control and fresh groups (both p<0.0001). Edema compared to initial weight at the time of procurement due to warm ischemia for 60 minutes at room temperature during simulation of the anhepatic phase of graft implantation (**Fig 4. H**) before the start of NMP was not different comparing CEPT and control groups (based on two-tailed two-sample student’s t-test with Welch’s correction). Weight changes at the end of 6 hours of NMP (**Fig 4. I**) compared to initial weight were also not different among all three groups (based on one-way ANOVA).

**Fig 4:**
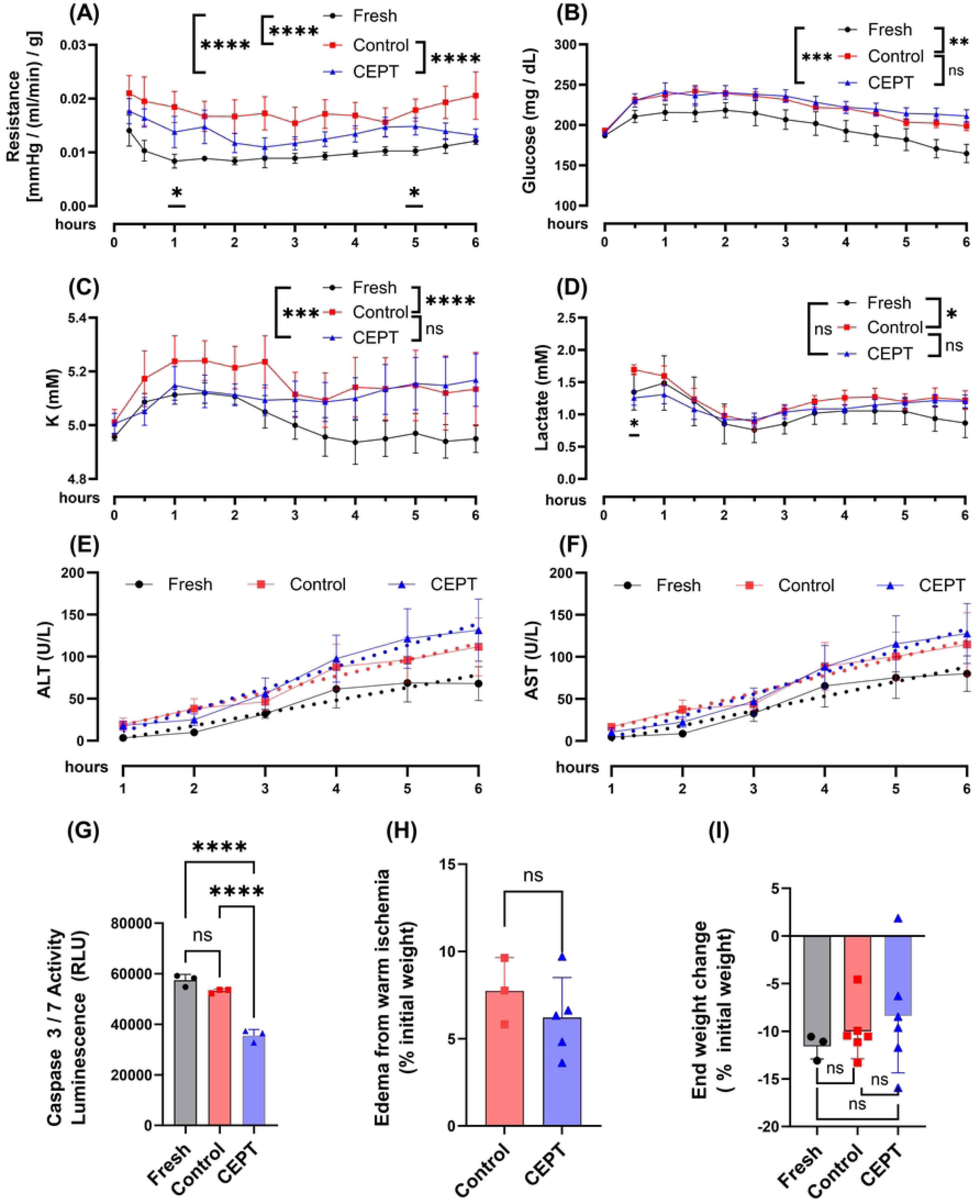
Injury markers during Normothermic Machine Perfusion (NMP): (A) Portal vascular resistance, (B) glucose (inflow), (C) K: potassium (inflow), (D) lactate (inflow), liver transaminase from inflow – (E) ALT and (F) AST, (G) Caspase 3/7 activity at the end of NMP, (H) edema from warm ischemia @ 21oC during the simulated anhepatic phase of graft implantation compared to initial weight at time of procurement, (I) end weight change after 6 hours of NMP compared to initial weight at the time of procurement. CEPT-treated livers had lower portal vascular resistance overall compared to the control group on One-way ANOVA (indicated by **** near groups legend), and at 1hr and 5hr, there was a significant difference between only the control and fresh group based on two-way ANOVA (indicated by * below line graph). Lactate was significantly different at 0.5 hr between the CEPT and control group based on two-way ANOVA (indicated by * below line graph). CEPT-treated livers had lower apoptosis marker indicated by lower Caspase 3/7 activity compared to the control group, but other injury markers were not different from the control group. Fresh (n=3): healthy control without 60 min warm ischemia time (WIT) at 21oC, control (n=6): with 60 min WIT @21oC but without CEPT, CEPT (n=6): experimental group with 60 min WIT @ 21oC and with CEPT; ANOVA with Tukey’s post-hoc test for multiple comparisons was used for all except (E), (F) & (H). For (E) & (F): simple linear regression with comparison of slopes with ANOVA and for (H): two-tailed two-sample student’s t-test with Welch’s correction; results are expressed as mean with SEM; pre-perfusion (time: 0) inflow glucose and K values before the livers were placed on the pump were excluded for analysis; ns: p > 0.05, * : p ≤ 0.05, ** : p ≤ 0.01, *** : p ≤ 0.001, **** : p ≤ 0.0001; ALT: Alanine transaminase, AST: Aspartate transaminase, K: Potassium.

### 3) Cholangiocellular function

Overall, the hourly rate of bile production (**Fig 5. A**) in the CEPT group was significantly higher than the control based on one-way ANOVA, and it was higher than the control group at 1 hr (0.05±0.002 vs. 0.038±0.004 ml/hr/g respectively, p = 0.038) based on two-way ANOVA. Still, both groups’ hourly bile production rates were not different from the fresh group liver. Bile pH delta, HCO_3_ delta, and K delta (**Fig 5. B-D**) were not different among the groups.

**Fig 5:**
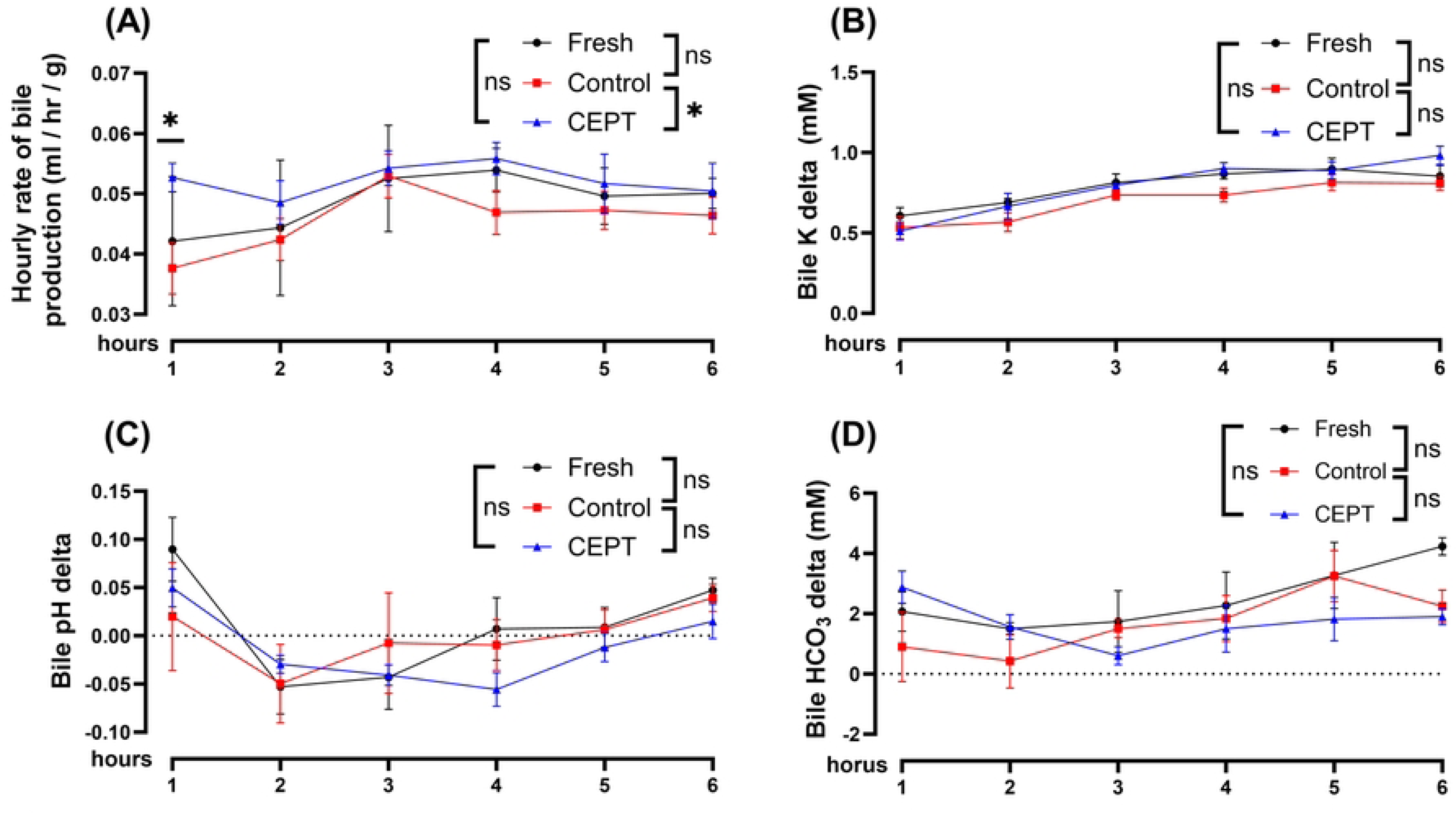
Cholangiocellular function during Normothermic Machine Perfusion (NMP): (A) Hourly rate of bile production (B) bile K delta (bile K - inflow perfusate K), (C) bile pH delta (bile pH - inflow perfusate pH), (D) bile HCO_3_ delta (bile HCO_3_ - inflow perfusate HCO_3_). Hourly rate of bile production was significantly higher overall on one-way ANOVA (indicated by * on legends) and was significantly higher during 1st hour of bile collection on two-way ANOVA (indicated by * on line graph) compared to the control groups. There was no significant difference among the groups for bile delta parameters. Fresh (n=3): healthy control without 60 min warm ischemia time (WIT) at 21oC, control (n=6): with 60 min WIT @21oC but without CEPT, CEPT (n=6): experimental group with 60 min WIT @ 21oC and with CEPT; ANOVA with Tukey’s post-hoc test for multiple comparisons; results are expressed as mean with SEM; ns: p > 0.05, * : p ≤ 0.05, ** : p ≤ 0.01, *** : p ≤ 0.001, **** : p ≤ 0.0001; HCO_3_: bicarbonate.

### 4) Histological analysis

Analysis of pathology slides using H&E staining (**Fig 6: A, C, E**) under light microscopy revealed no significant differences among the groups in terms of tissue necrosis and interstitial edema. The portal triad (indicated by the structure around the central vein with a star (*)) was well preserved without cellular and interstitial edema. TUNEL staining (**Fig 6: B, D, F**) showed occasional TUNEL-positive nuclei in high power fields (200x magnification) and were similar among the groups.

**Fig 6:**
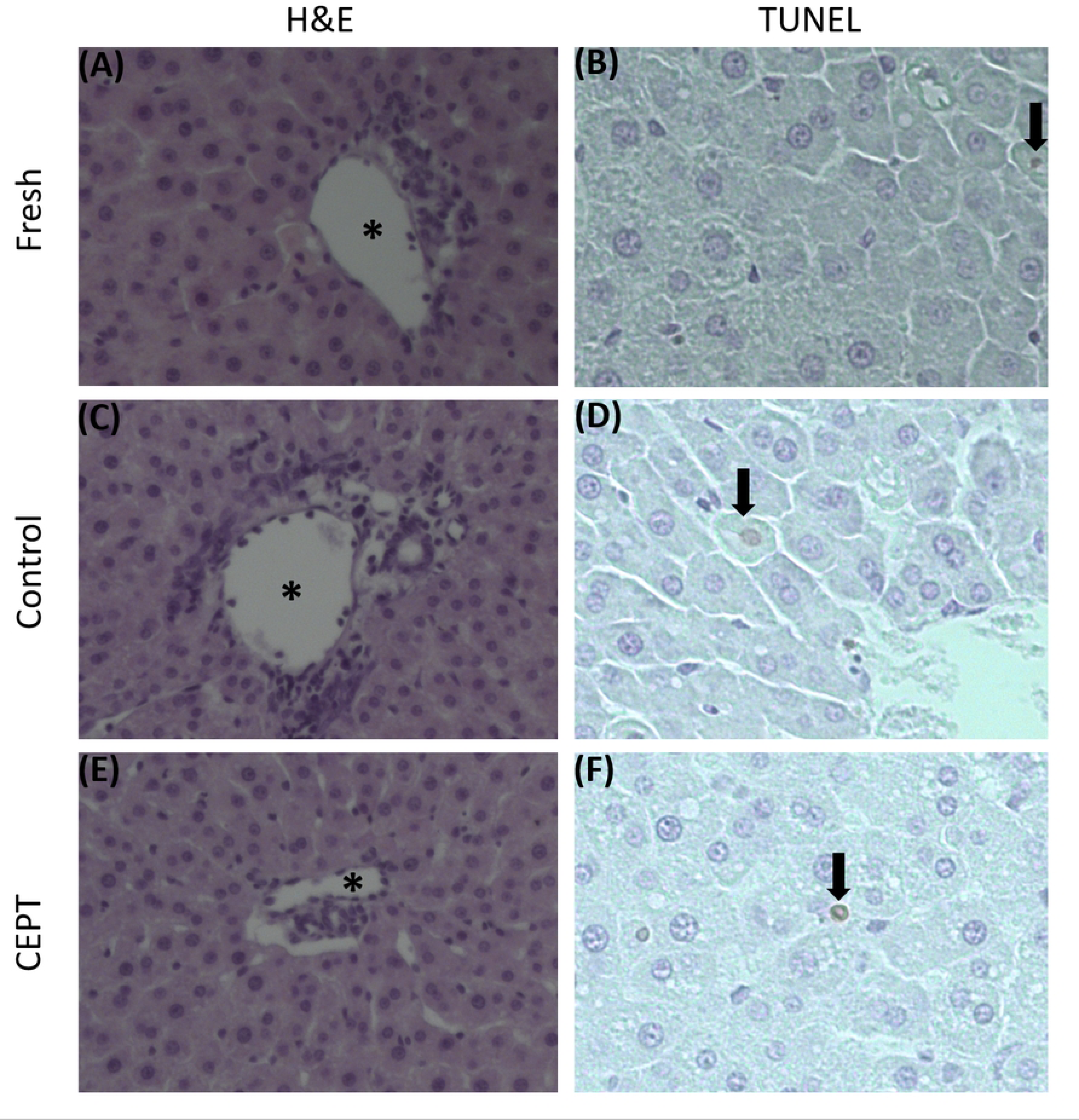
Histology with H&E and TUNEL staining and light microscopy of the liver biopsy taken at the end of 6 hours of NMP: H&E staining at 100x magnification- (A) fresh group, (C) control group, (E) CEPT group; TUNEL staining at 200x magnification- (B) fresh group, (D) control group, (F) CEPT group. Light microscopy images using H&E staining did not show a significant difference among the groups, and TUNEL staining used for visualizing DNA fragmentation for identifying apoptotic cells showed occasional TUNEL positive nuclei and were similar among the groups; star (*): portal vein with focus on portal triad, black arrow represents TUNEL stained nuclei. H&E: Hematoxylin & Eosin, TUNEL: Terminal deoxynucleotidyl transferase dUTP nick end labeling.

### 5) Storage solution and Effluent drainage analysis

Storage solution (**S2 Fig**) in which the control and CEPT groups grafts were maintained at room temperature for inducing warm ischemia for simulating anhepatic phase and effluent drainage (**S3 Fig**) collected from IVC just before the start of NMP were not different among the groups in terms of biochemistry panels: pH, Lactate, Glucose, K, and transaminase: ALT & AST. In addition, AST and ALT were not detectable in the storage solution and, therefore, were not shown in the graph.

## Discussion

Our study simulates injury from warm ischemia during the anhepatic phase of graft implantation in the recipient, as well as simulates 6 hours of the post-reperfusion phase of the transplantation process using an NMP pump with an exploration of the possibility of drug intervention for minimizing IRI. This study suggests that flushing with CEPT and treatment with CEPT post-reperfusion of the graft in a transplant simulation model reduces damage associated with reperfusion, as demonstrated by increased O2 uptake, pH stability (Fig 3. A & B), higher hourly rate of bile production (**Fig 5. A**) and lower portal vascular resistance (**Fig 4. A**), lower caspase 3/7 activity (**Fig 4. G**), and initial lower lactate level at half an hour timepoint (**Fig 4. D**).

Improved portal vascular resistance in the CEPT group could result from decreased endothelial injury from induction of warm ischemia, suggesting a lowering of endothelial cell swelling in the liver sinusoids and ultimately keeping the capillary vasculature dilated. Low lactate levels and effective lactate clearance both in perfusate and bile have been used for viability assessment [20], and CEPT-treated liver recovered faster, showing lower lactate at the half an hour timepoint of perfusion. Hyperglycemia, non-responsiveness to insulin, is associated with aggravated IRI. Therefore, elevated glucose levels in the perfusion media may serve as an additional injury marker [21]. Both the control and CEPT groups manifest higher overall glucose levels when compared to the fresh group, indicating higher IRI from the induction of warm ischemia of 60 minutes at room temperature. The linear trend with time of transaminase level can be explained by the stress of the machine perfusion system and the lack of dialysis circuits in our perfusion system. Both CEPT and control groups experienced similar levels of edema from the induction of 60 minutes of warm ischemia to simulate the anhepatic phase.

Similarly, weight loss in all groups could be caused by recovery of cellular swelling [22] that occurred during the liver procurement phase and the 60 minutes of WIT phase in the CEPT and control groups. RL solution instead of the University of Wisconsin-Cold Storage Solution (UW-CSS) was used for flushing the graft during procurement since our study did not involve static cold storage (SCS), which is unavoidable in clinical practice because of the need for graft transport. This was done to remove the cold ischemic time (CIT) of SCS as a confounding variable. It is worth noting that the RL solution does not contain colloids such as hydroxyethyl starch, present in UW-CSS for balancing the hydrostatic pressure in the sinusoids during procurement flush, and this could be responsible for causing more edema at the time of procurement, which can result in falsely elevated initial weight after procurement. This can be the second reason for weight loss seen among all three groups at the end of 6 hours of NMP.

Higher oxygen uptake and stable pH in the CEPT group could be due to a greater number of healthy hepatocytes compared with the control group. The bile volume produced has been used as a biomarker of hepatic viability during ex vivo machine perfusion [23], and we observed that the CEPT group’s hourly bile production is higher compared to the control group. This difference is particularly noticeable in the first hour, which may be due to the protective effect of CEPT on cholangiocytes during flushing and WIT. Cholangiocytes function to secrete bicarbonate, resulting in bile with an alkaline pH (>7.45) [24]. The injured biliary tree cannot secrete sufficient HCO3, which results in reduced basic bile. Therefore, biliary delta parameters calculated from the difference between bile and inflow perfusate, like pH delta, HCO3 delta, and K delta, have been used as biomarkers of biliary duct injury [24, 25]. This study could not show differences among the groups using these bile delta parameters, possibly due to a relatively low level of warm ischemic damage at room temperature for 60 minutes.

There are several limitations from the perfusate used in this study. First, the perfusate does not have an oxygen carrier. Without an oxygen carrier, the delivery of oxygen to the graft is limited, and all metabolic parameters presented in this study are slightly lower than what would have been possible with an added oxygen carrier because a graft can only extract a fraction of the oxygen delivered [26]. The addition of an oxygen carrier in the perfusate might have magnified the difference among the groups related to various parameters, helping to reach the statistical significance level. Furthermore, a comparatively large volume of perfusate compared to graft weight causes the dilution of injury markers like lactate, K, and ALT/AST, making it challenging to observe significant differences. Our study relies on a simulation of 60 minutes of the anhepatic phase of graft implantation and 6 hours of post-reperfusion in the recipient on a pump without using whole blood as perfusate. Due to the lack of whole blood in perfusate, it fails to reveal a high level of IRI from damage caused by innate and acquired immunity like WBCs activation, complement pathway activation, and effects of various cytokines [2]. Furthermore, the short duration of perfusion of 6 hours lacks long-term follow-up.

## Conclusion

CEPT targets various apoptotic pathways [11] and minimizes IRI from the gradual rewarming during the simulated anhepatic phase of graft implantation and enhances graft recovery post-reperfusion, as indicated by significant improvement in hepatocellular function and injury markers. Both the liver graft and patient survival are inferior among recipients of DCD livers compared to DBD [27], and the main factor limiting the utilization of marginal livers from DCD is the subsequent development of ischemic-type biliary lesion (ITBL) [28]. Since CEPT is found to be beneficial for the graft to handle the stress of mild warm ischemia at room temperature, we can try exploring the possibility of recovering DCD liver grafts, which have suffered severe warm ischemic damage due to remaining at a body temperature of 37^0^C without circulation. Further experiments with the addition of CEPT to individual flush, storage, and perfusate solutions are needed to stratify its protective effect. Treatment of brain-dead donor (DBD) for upstream IRI minimization and recipient animals during and after graft implantation for downstream IRI minimization using CEPT is to be explored in future in-vivo studies.

## Acknowledgments

All illustrations were created with BioRender.com (Toronto, ON, Canada).

## Supporting information

**S1 Fig: Graphical abstract**

**S2 Fig: Biochemistry analysis of storage Ringer’s Lactate (RL) solution, in which procured liver graft was stored for 60 min at room temperature (21°C) to induce warm ischemic damage to simulate the sewing phase (anhepatic phase) of the graft implantation in the recipient**. (A) pH, (B) lactate, (C) K: potassium, (D) glucose. Storage solution biochemistry analysis did not show a significant difference. The solid line represents the lower values of pH and glucose that the diagnostic machine can measure, and the dotted line represents the normal concentration of lactate and K in RL. Control (n=3): with 60 min WIT @21°C but without CEPT, CEPT (n=4): experimental group with 60 min WIT @ 21°C and with CEPT; two-tailed two-sample student’s t-test with Welch’s correction; results are expressed as mean with SEM.

**S3 Fig: Effluent drainage analysis collected from inferior vena cava (IVC) after flushing with 3ml of Ringer’s Lactate (RL) just before the start of Normothermic Machine Perfusion (NMP).** (A) pH, (B) glucose, (C) ALT, (D) lactate, (E) K: potassium, (F) AST. Effluent drainage biochemistry analysis and transaminase did not show a significant difference. Solid line represents the lower values of pH and glucose that the diagnostic machine can measure, and the dotted line represents the normal concentration of lactate and K in RL. Also, the dotted lines in ALT and AST represent the upper level of physiological transaminase value. Fresh (n=3): healthy control without 60 min warm ischemia time (WIT) at 21°C, control (n=3): with 60 min WIT @21°C but without CEPT, CEPT (n=4): experimental group with 60 min WIT @ 21°C and with CEPT; ANOVA with Tukey’s post-hoc test; results are expressed as mean with SEM; ALT: Alanine transaminase, AST: Aspartate transaminase.

**S4 Data**

## Notes

### Competing Interest Statement

Some authors declare competing interests. MT, SNT, HY, and KU have patent applications for apoptotic inhibitors, and MT, SNT, and KU have a financial interest in and serve on the Scientific Advisory Board for Sylvatica Biotech Inc., a company focused on developing high subzero organ preservation technology. MGH and MGB manage competing interests for MGH investigators through their conflict-of-interest policies.

